# Investigating the efficacy of triple artemisinin-based combination therapies (TACTs) for treating *Plasmodium falciparum* malaria patients using mathematical modelling

**DOI:** 10.1101/331850

**Authors:** Saber Dini, Sophie Zaloumis, Pengxing Cao, Ric N Price, Freya J.I. Fowkes, James M McCaw, Julie A Simpson

**Author notes:** Address correspondence to Julie A. Simpson,.

## Abstract

The first line treatment for uncomplicated falciparum malaria is artemisinin-based combination therapy (ACT), which consists of an artemisinin derivative co-administered with a longer acting partner drug. However, the spread of *Plasmodium falciparum* resistant to both artemisinin and its partner drugs poses a major global threat to malaria control activities. Novel strategies are needed to retard and reverse the spread of these resistant parasites. One such strategy is triple artemisinin-based combination therapy (TACT). We developed a mechanistic within-host mathematical model to investigate the efficacy of a TACT (dihydroartemisinin-piperaquine-mefloquine - DHA-PQ-MQ), for use in South-East Asia, where DHA and PQ resistance are now increasingly prevalent. Comprehensive model simulations were used to explore the degree to which the underlying resistance influences the parasitological outcomes. The effect of MQ dosing on the efficacy of TACT was quantified at varying degrees of DHA and PQ resistance. To incorporate interactions between drugs, a novel model is presented for the combined effect of DHA-PQ-MQ, which illustrates how the interactions can influence treatment efficacy. When combined with a standard regimen of DHA and PQ, the administration of three 8.3 mg/kg doses of MQ was sufficient to achieve parasitological efficacy greater than that currently recommended by WHO guidelines.

## Introduction

Over the last decade, significant gains have been made towards the control and elimination of malaria. Despite this progress, almost half a billion people still die from malaria each year. Disturbingly, in 2016 there were five million more cases of malaria than the previous year (2017 WHO report (1)), emphasising the fragile nature of malaria control. Early diagnosis and treatment with highly effective antimalarial drug regimens remains central to all national malaria control activities. Artemisinin-based combination therapies (ACTs) are the first line therapy in almost all malaria endemic countries, due to their high efficacy, tolerability and ability to reduce ongoing transmission of the parasite. ACTs are comprised of two components: an artemisinin derivative and a partner drug. The artemisinin derivative has a high antimalarial potency, killing a large proportion of parasites, however, these compounds are rapidly eliminated, leaving a residual parasite population that, if left untreated, will likely recrudesce. A slowly eliminated, partner drug, is required to provide a sustained antimalarial activity, capable of killing the remaining parasites (2).

ACTs have remained highly efficacious for almost two decades, but are now under threat from the emergence of drug resistant parasites (2, 3). In 2009, a high proportion of patients with markedly delayed parasite clearance were reported from the western region of Cambodia, and this was confirmed as being attributable to artemisinin resistance (3). These parasites have now spread across the Greater Mekong Region (4, 5). Delayed parasite clearance and higher gametocyte carriage, due to the artemisinin derivative resistance, drive selection of resistance to the partner drug (6), and in South-East Asia, this has resulted in declining efficacy of all the ACTs currently recommended by WHO (7). In some parts of the Greater Mekong Region, the spread of highly drug resistant parasites poses a major threat to malaria control activities. The emergence of an untreatable *P. falciparum* will result in an inevitable rise in malaria incidence, epidemics and associated morbidity and mortality.

The development of alternative strategies is crucial to ensuring the ongoing success of malaria control efforts. Triple Artemisinin-based Combination Therapy (TACT) is a novel strategy by which a new partner drug is added to an established ACT. TACT has the potential to prevent the emergence of a *de novo* resistance as well as rescuing a regimen in which one of the ACT components is already failing. Two antimalarial clinical trials are underway to determine the efficacy of TACT for uncomplicated falciparum malaria: Artemether-Lumefantrine plus Amodiaquine (AL-AQ) and Dihydroartemisinin-Piperaquine plus Mefloquine (DHA-PQ-MQ). These are being compared to the standard ACTs (AL and DHA-PQ) alone (see trial number NCT02453308 in clinicaltrials.gov).

In this work, we developed a within-host mathematical model (8) to explore the efficacy of TACTs, with a particular focus on DHA-PQ-MQ, since DHA-PQ is widely administered in South-East Asia, and is currently associated with very high failure rates in some regions (9, 10, 11). The model accommodates a high level of biological details, such as drug-drug interaction (12, 13), stage-specificity of parasite killing (14, 15, 16, 17) and between-patient and between-isolate variability (17). We used the model to simulate different levels of resistance and quantify the degree to which this compromises the efficacy of TACT. The optimal MQ dosing regimen was determined for various degrees of resistance to DHA-PQ.

## Results

Simulated drug concentrations and parasite burden are shown in Fig. 1. The median concentration of the drugs (lines) along with the between-subject variabilities (the shaded regions show the area between the 2.5% and 97.5% percentiles) are presented in Fig. 1a, and the parasitaemia of 100 randomly selected patients in Fig. 1b. Fig. 1c presents the Kaplan-Meier estimation of the probability of cure, along with the 95% confidence intervals illustrated by the shaded region.

**Figure 1:**
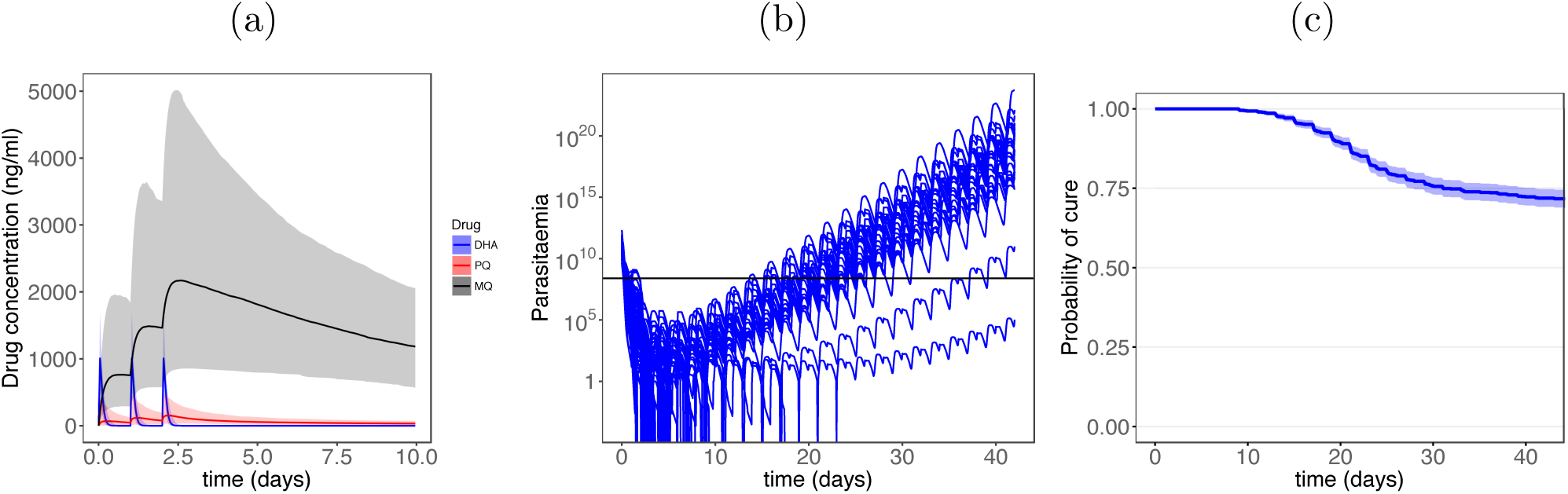
Model simulation. a) PK model results; the concentrations of DHA (blue), PQ (red), MQ (black) are depicted. The shaded regions show the area between the 2.5th and 97.5th percentiles of the results generated for 1000 patients. b) PD model results for 100 randomly selected patients; the horizontal line shows the microscopic level of detection of parasites. c) Kaplan-Meier estimation of the probability of survival over 42 days of follow-up.

Parasite resistance to antimalarial drugs can manifest in a couple of different ways that affect the killing profile of a drug (see the concentration-effect curves in Fig. 2 (18)). These include i) increasing *EC*_50_ (the red curve), ii) reducing the size of the killing window in the intra-erythrocytic parasite life-cycle, and iii) decreasing *γ* and/or *E*_*max*_ (the blue curve). The degree of resistance was modelled initially by varying *EC*_50_ of PQ, in scenarios where the parasites are sensitive or resistant to DHA. The influence of other manifestations of resistance on TACT efficacy are outlined in Supplementary Material 1.

**Figure 2:**
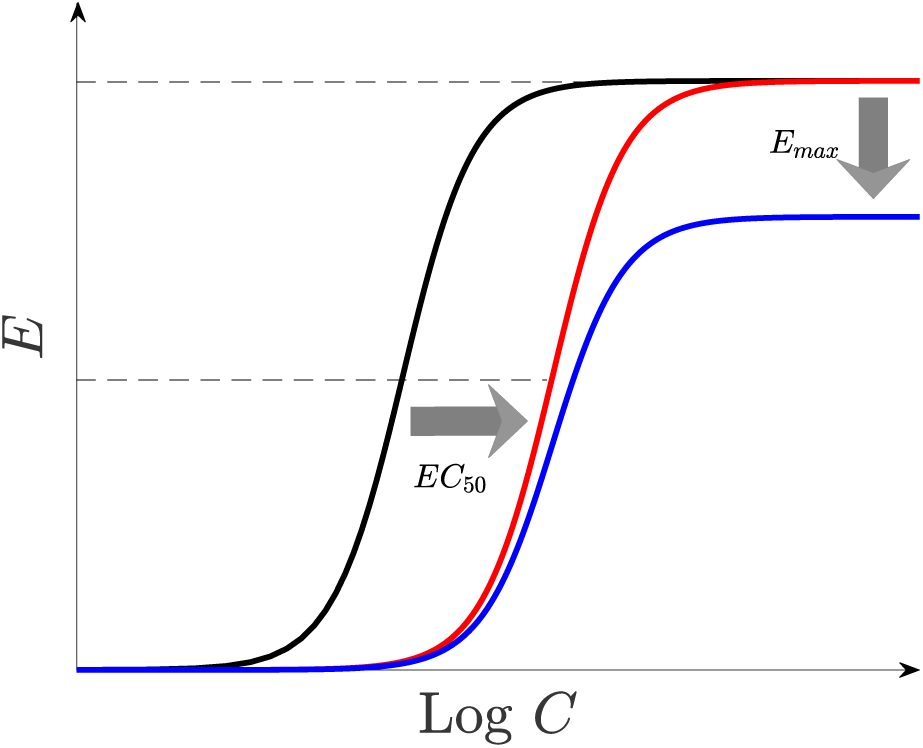
Resistance manifestations. Resistance of parasites to drugs, modelled by relevant alterations of the parameters of the model. A concentration-effect profile of susceptible parasites (black) can be right-shifted, *i.e. EC*_50_ increases (red), and/or the maximum killing effect, *E*_*max*_, decreases (blue).

The level of resistance and the resultant risk of treatment failure varies with geographical region. Table 1 demonstrates a large variation in DHA-PQ efficacy in different regions across South-East Asia (19). The risk of failures in Aoral and Chi Kraeng in Cambodia are 51.9% and 62.5% treatment failures, respectively, whereas in Siem Pang it is only 8.3%. Similar large variations in the probability of treatment failure are observed in Vietnam. According to the WHO treatment guidelines, when the risk of failure exceeds 10%, a treatment is considered suboptimal, and steps should be taken to change the policy to a more efficacious antimalarial regimen.

**Table 1:** The Kaplan-Meier estimation of the probabilities of cure on day 42 of follow-up in some regions in South-East Asia where DHA-PQ is the first-line treatment for malaria (19).

|  | Geographical Region |  |  |  |  |
| --- | --- | --- | --- | --- | --- |
|  | Siem Pang | Binh Phuoc | Bu Gia Map | Aoral | Chi Kraeng |
| Probability of cure | 0.92 | 0.74 | 0.67 | 0.48 | 0.38 |
| Number of patients | 60 | 40 | 40 | 53 | 40 |
| Year | 2015 | 2015 | 2015 | 2015 | 2014 |
| Country | Cambodia | Viet Nam | Viet Nam | Cambodia | Cambodia |

### Sensitivity to DHA

In the first investigated scenario, the parasites were assumed sensitive to DHA (sampling interval of *EC*_50,*D*_ was limited to (0, 10] ng/ml), and resistance level to PQ was varied. Fig. 3a shows how the probability of cure at day 42 of follow-up varies with *EC*_50_ of PQ over the deciles of (11, 94] ng/ml. The top labels in this figure show the geographical regions in South-East Asia (Table 1) that have observed DHA-PQ day 42 cure rates equal to the corresponding simulated values (19).

**Figure 3:**
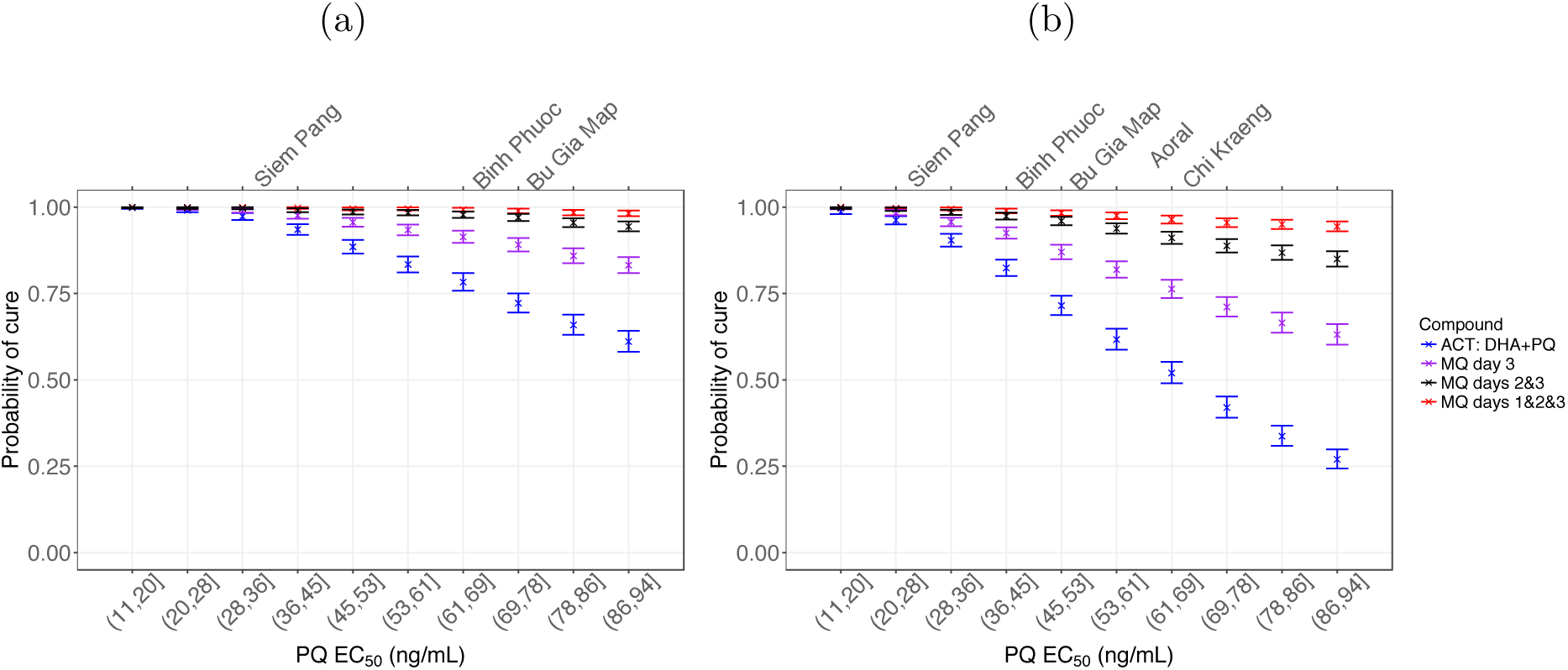
The probability of cure on day 42 of follow-up when *EC*_50_ of PQ varies over the deciles of [11 94]. (a) Sensitivity and (b) resistance to DHA. Blue: ACT treatment — dosing regimens of PQ and DHA are 18.0 mg/kg and 4.0 mg/kg, respectively, on days 1, 2 and 3. Purple: a single dose of 8.3 mg/kg of MQ is added on day 3. Black: two 8.3 mg/kg doses of MQ on days 2 and 3 are added. Red: three 8.3 mg/kg doses of MQ are added on days 1, 2,3. The top labels show the geographical regions in South-East Asia (Table 1) that have observed DHA-PQ cure rates equal to the corresponding simulated values. Error bars show the 95% confidence intervals of Kaplan-Meier analysis.

The probability of cure declines as *EC*_50_ of PQ increases. Without MQ, the probability of cure with DHA-PQ is below 90%, over *EC*_50,*P*_ ∈ (36, 94], which includes Binh Phuoc and Bu Gia Map. This scenario was unable to produce the probabilities of cure observed in all of the geographical regions, shown in Table 1.

The addition of a single 8.3 mg/kg dose of MQ on day 3 significantly raised the probability of cure. Day 3 administration of MQ was chosen, since at this time patients are clinically better, the drug is better tolerated and bioavailability is higher (20). The improvement in efficacy with a single dose of MQ was insufficient to ensure successful treatment in Bu Gia Map. In this region, a second dose at day 2 was required. When the parasites are sensitive to DHA, but resistant to PQ, two doses of MQ on days 2 and 3 were sufficient to achieve cure in all locations. Administration of three doses of MQ did not provide significant benefit over a two dose MQ regimen, although might be used to guarantee the success of the TACT.

### Resistance to DHA

Concurrent resistance to DHA and PQ is now documented in Cambodia and Vietnam (9). To simulate a high level of DHA resistance, we set *EC*_50,*D*_ ∈ (50, 100] ng/ml, and varied the intensity of resistance to PQ, *EC*_50,*P*_ ∈ (11, 94] ng/ml. Using the same dosing regimens as those in Fig. 3a, resistance to DHA leads to a significant decline in the efficacy of DHA-PQ, as shown in Fig. 3b. When combined with a single 8.3 mg/kg dose of MQ, the efficacy of the TACT was improved significantly, but except for Siem Pang, it was clearly not sufficient.

Administration of MQ on days 2 and 3 provided sufficient efficacy in Binh Phuoc and Bu Gia Map, but was still insufficient for Aoral and Chi Kraeng. An additional dose of MQ was needed on day 1 to obtain a successful treatment in all of the regions. Note that administering 8.3 mg/kg of MQ over three days (25 mg/kg in total) is currently the recommended dosing regimen by WHO for the ACT of MQ plus artesunate (18).

### The influence of the antagonistic PQ-MQ interaction

The effect of the PQ-MQ interaction parameter, *α*, on the probability of cure was then investigated. Fig. 4a shows the combined killing effect of the drugs, *E*, over time for a selected patient with two different values of the interaction parameter: *α* = 3.3 (antagonism) and *α* = 1 (zero-interaction); the other parameters were kept constant. The killing effect for *α* = 1 (solid line) is significantly higher than that for *α* = 3.3 (dashed line), indicating the extent to which the drugs can nullify each other’s effect, and the loss in the overall efficacy of TACT.

**Figure 4:**
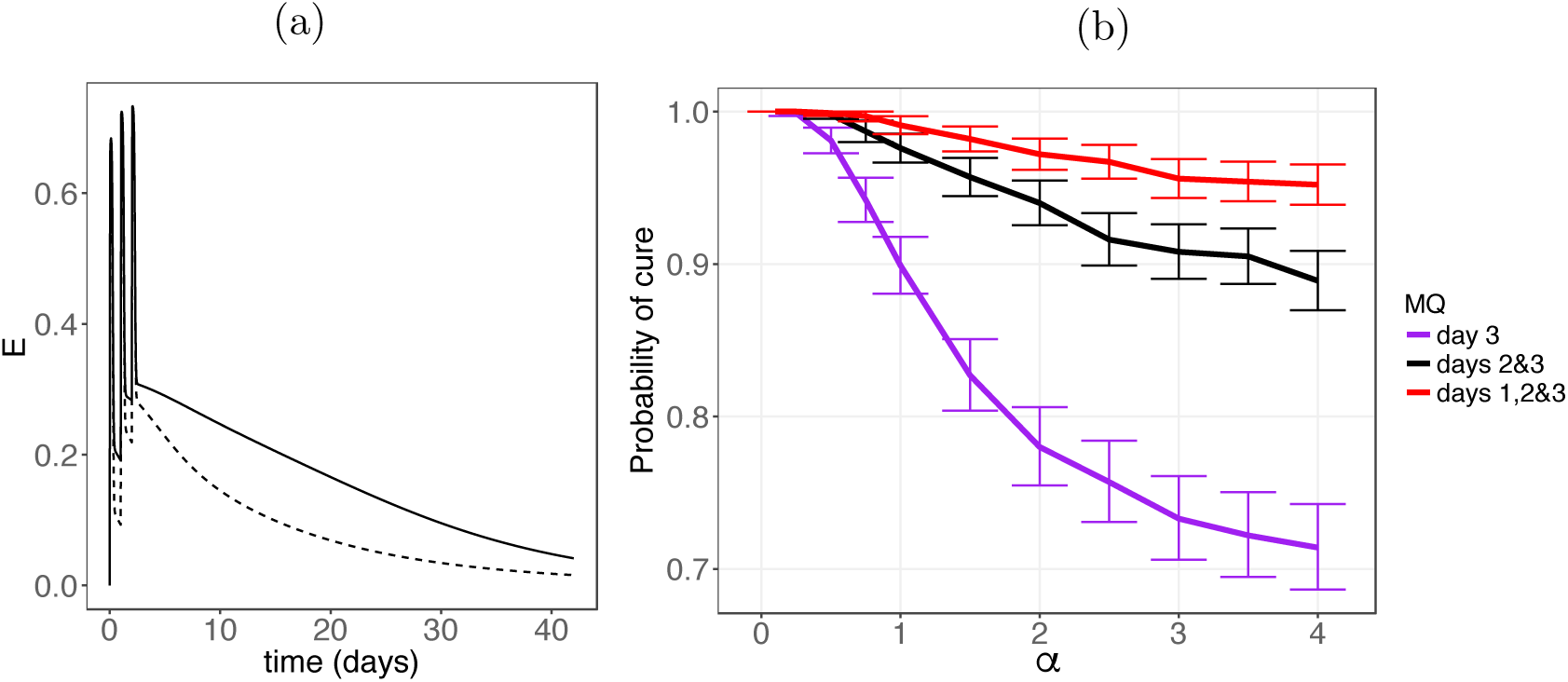
The influence of antagonism between PQ and MQ on the efficacy of the TACT. (a) Dashed and solid lines represent combined killing effect, *E*, for *α* = 3.3 and *α* = 1, respectively. (b) The probability of cure on day 42 of follow-up versus the interaction parameter, *α*, when the resistance level corresponding to Chi Kraeng is considered, *i.e. EC*_50,PQ_ ∈ (69, 78]; resistance to DHA is assumed. Different values of interaction parameter, *α*, produce synergism (0 < *α* < 1), zero-interaction (*α* = 1) and antagonism (1 < *α* < ∞) in the combined e*α*ect of PQ-MQ. The interpretation of the colors is explained in the caption of Fig. 3.

The effect of the interaction parameter, *α*, on the efficacy was further assessed by restricting the resistance level to that corresponding to Chi Kraeng (*EC*_50,*P*_ ∈ (69, 78]) and estimating the probability of cure for different values of *α*; DHA resistance is also assumed. The results demonstrated a significant difference between the probabilities of cure at different values of *α*; Fig. 4b. For example, when *α* < 1 (synergism), one dose of MQ was enough to provide 90% efficacy. In contrast, when *α* > 1 (antagonism) the probability of cure fell well below 90%. Similarly, the probability of cure declined with increasing *α* (*i.e.* antagonism intensification) for MQ administration on days 2 and 3 and on days 1, 2 and 3. Of note, the three 8.3 mg/kg doses of MQ achieved greater than 90% efficacy at all values of *α*, even at levels indicative of very strong antagonism. This highlights the robustness of this dosing regimen in producing a successful treatment. The antagonism between PQ and MQ had an important impact on the efficacy of the TACT, and neglecting this may result in an underestimation of the dose of MQ required for successful treatment across different regions.

The effect of other manifestations of resistance on the efficacy of the TACT are illustrated in Supplementary Material 1. Fig. S1 presents the probability of survival at different levels of resistance produced by varying maximum killing effect of PQ, *E*_*max,P*_. Similar to the case where *EC*_50_ was the manifestation of resistance, shown in Fig. 3, the results indicate that the three 8.3 mg/kg doses of MQ are sufficient to provide the desirable probability of cure at every level of resistance. The outcomes were consistent when the killing window of PQ was shortened, as shown in Fig. S2. However, the probability of cure became extremely low, when resistance was high. Nevertheless, three 8.3 mg/kg doses of MQ overcame high levels of resistance and ensured high probability of cure.

## Discussion

We have presented a novel mathematical model to investigate the efficacy of different regimens for triple artemisinin combination therapies (TACTs). Our analysis focused on DHA-PQ-MQ, since DHA-PQ is a widely used ACT in South-East Asia with declining efficacy in several locations (9, 10, 11). The addition of MQ to DHA-PQ has potential to improve treatment, since the ACT of artesunate-MQ retains high efficacy, following its reintroduction as a first-line treatment in Cambodia (7). Our results suggest that a single dose of MQ can improve the treatment efficacy of DHA-PQ significantly, and would be an appropriate regimen for regions such as Siem Pang in Cambodia and Binh Phuoc in Viet Nam. However, it is likely to be insufficient in regions where there is pre-existing high grade resistance to PQ, such as Bu Gia Map in Viet Nam and Aoral and Chi Kraeng in Cambodia. The addition of two doses of MQ would be beneficial, but efficacy would still be compromised in the regions where there was high level of resistance to both PQ and DHA, such as Chi Kraeng. To achieve a cure rate of greater than 90%, as recommended by the WHO, three doses of MQ (8.3 mg/kg) need to be administered in conjunction with the standard three days regimen of DHA-PQ. Such a dose of MQ, has already been shown to be well tolerated and safe, and is recommended in the WHO treatment guidelines (18).

Our model enabled us to simulate the PK and PD following TACT administration to patients with malaria, and provided important insights into the way in which the underlying mechanisms of drug action affect treatment efficacy. By taking account of between-patient and between-isolate variability, we were able to explore treatment efficacy across a wide range of different scenarios reflecting varying parasite resistance to the different drug components. The results showed similar trends for different resistance manifestations, confirming the robustness of the proposed dosing regimen of DHA-PQ-MQ.

We have proposed a novel empirical model to accommodate the effect of the combined drugs, assuming that PQ and MQ (both quinoline compounds) have similar modes of action, which differs from that of DHA (an endoperoxide compound). The killing effects of PQ-MQ and DHA were therefore assumed to be independent. This justified using a combination of Bliss independence and Loewe additivity to define the combined effect of the whole compound (see Eqn. (1)).

To facilitate the dissemination of our model and assist clinical researchers to investigate how different PK and PD parameters and dosing schemes influence parasitological outcomes, we have produced an online application that allows varying the values of parameters and simulating the model: appTACT. By predicting the fate of malaria infection in patients, the online application can provide a means for *e.g.* estimating the required sample size of an antimalarial clinical efficacy study.

Our mathematical model can be used to guide the development of suitable TACT regimens for investigation in clinical trials. Determining dosing regimens that are robust to a wide range of scenarios helps rationalize the logistical and financial challenges of phase 2 and 3 clinical trials. Further improvements in the model can be made to increase its fidelity to the underlying biology. For instance, by consideration of the artemisinins PK (e.g. bioavailability) dependence on parasite density (21) and different bioavailability of MQ at different administered days (20). The PD model can also be improved by incorporating more complexities underlying drug action, such as the dependence of killing effect on the timespan parasites are exposed to drugs (22, 23, 24, 25). However, in this initial analysis, we aimed to focus on the generality of the model and leave these modifications for future studies. Although we did not explore the degree to which the efficacy of TACT influenced other aspects of malaria control, such as the transmissibility of the parasite, this certainly warrants further investigation, since a more comprehensive perspective will be needed on the suitability of deploying TACT in areas of high drug resistance.

## Materials and Methods

### Mathematical Model

The pharmacokinetic-pharmacodynamic (PK-PD) model presented in Zaloumis et al. 2012 (17) was used to model the dynamics of drug concentrations and parasite burden within an individual. In brief, this model describes the time-evolution of the number of parasites in the body, N, by the following difference equations:

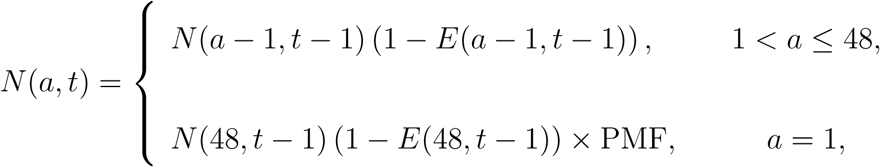

where *a* is the parasites’ age, taking only integer values over [1 48], *t* is time and PMF is the parasite multiplication factor, which represents the number of merozoites released into blood by a shizont at the end of its lifecycle. *E(a, t)* is the combined killing effect of the drugs, and has been modified from that presented in Zaloumis et al. 2012 to account for three drugs and accommodate drug interactions. The combined killing effect of the drugs is between 0 and 1, and dependent upon the age of parasites during [*t, t* + 1). The number of detectable parasites circulating in the blood, *M*(*t*), is determined by

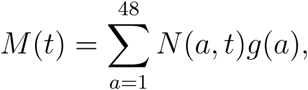

where *g(a)* accounts for the reduction in the number of circulating parasites in the blood due to sequestration, estimated to be

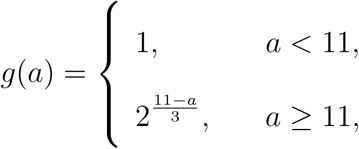

where we assumed sequestration begins at age 11 and intensifies with age (16, 26). In the ensuing section, we explain the details of modelling the combined effect of the drugs, *E*.

The PK models for the three drugs considered, DHA, PQ and MQ, are well characterized; one-compartment models were used for DHA and MQ and a two-compartment model for PQ (27, 28, 29). The PK parameter values are drawn from the literature and provided in Table 2.

**Table 2:**
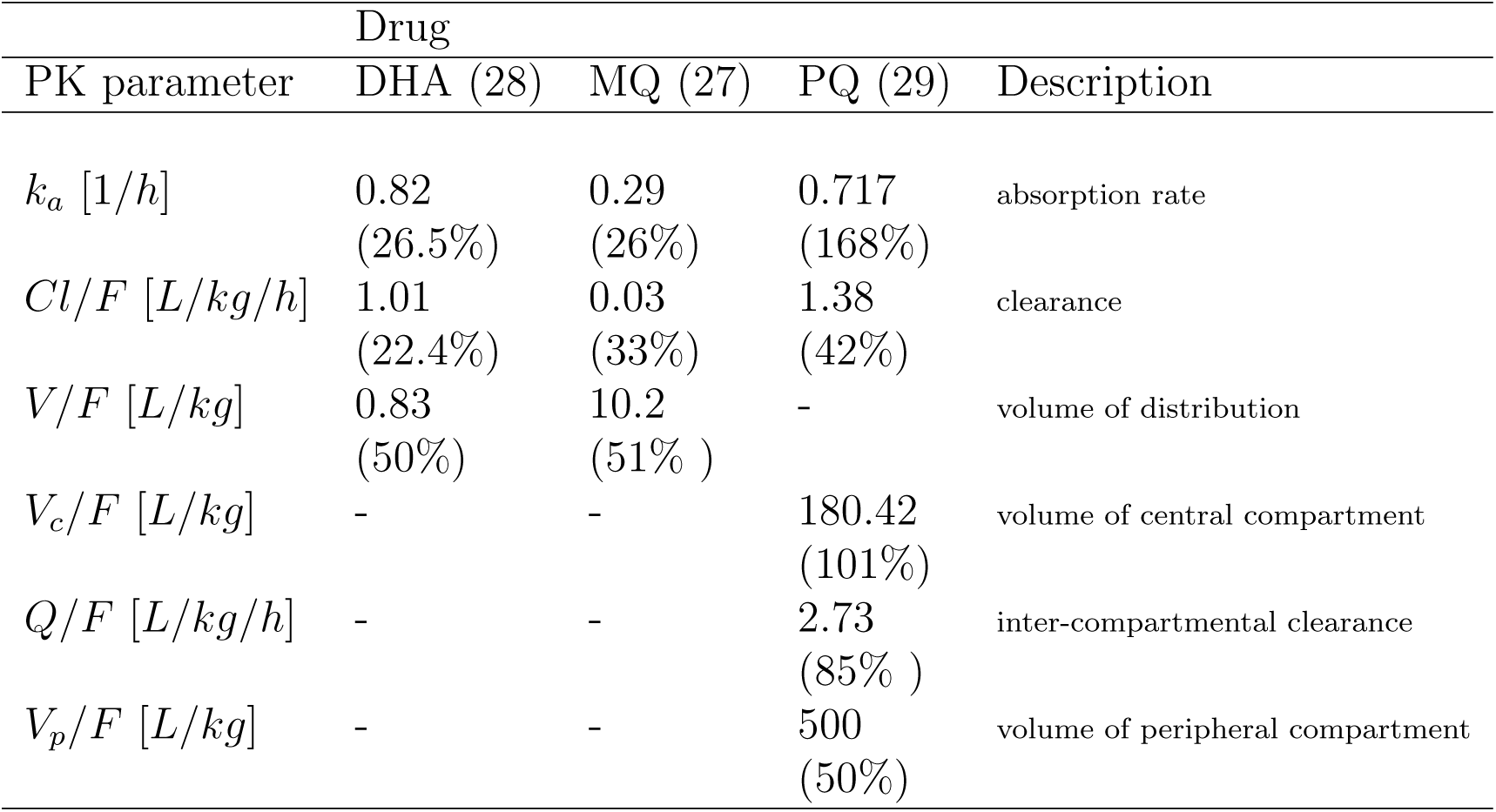
Parameter values of the pharmacokinetic model. The mean values are shown along with the between-patient variabilities (presented as % coefficient of variation) in brackets.

### Combined killing effect of the drugs

The combined killing effect of the drugs is modulated by the manner in which they interact with each other. Synergistic interaction between drugs produces a stronger combined effect compared to the case where they do not interact, *i.e.* zero-interaction (also known as pure additivity). Conversely, antagonistic drug-drug interactions can nullify their additive effect, and produce a lower combined effect than that for the zero-interaction case. Therefore, to model the combined effect, *E*, we must first identify how the drugs interact.

An empirical approach was taken, modelling zero-interaction as the reference (null) model (30, 31, 32), since the mechanisms underlying the killing effects are complex and not completely understood (33). Among the existing empirical approaches of modelling zero-interaction, two are more prominent and widely used: *Loewe additivity* (34) and *Bliss independence* (35). Loewe additivity is suggested to be a suitable concept for zero-interaction when non-interacting drugs have similar modes of action, however, when the drugs are believed to act independently, Bliss independence is more appropriate (31, 32).

It has been suggested that MQ and PQ kill parasites through a similar mechanism, involving the disruption of haem detoxification in the parasite vacuole (33, 36, 37). DHA has a different mode of action, which involves the generation of free radicals and reactive intermediates that target various proteins of parasites (36, 38, 39). The PK and PD interactions of DHA with PQ and MQ appear to be negligible (13).

The independent mechanisms of action of DHA and PQ-MQ justifies using the Bliss independence concept for modelling the combined killing effect, *E*, given by

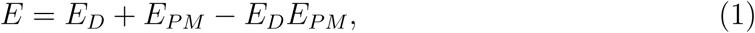

where *E*_*D*_ is the killing effect of DHA and *E*_*PM*_ is the combined effect of PQ and MQ. We assume Michaelis-Menten kinetics for *E*_*D*_:

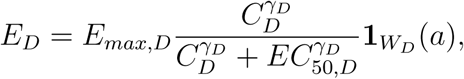

where *E*_*max,D*_ is the maximum killing effect of DHA; *C*_*D*_ is DHA concentration; *EC*_50,*D*_ is the concentration at which 50% of the maximum killing effect is obtained; *γ*_*D*_ is the sigmoidicity (also known as slope) of the concentration-effect curve; **1**_*W*_*(a)* is an indicator function, used to implement the age-specific killing of drugs, defined by

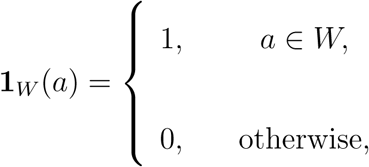

where *W* is the age window (interval) where the antimalarial drugs are able to kill the parasites; *W*_D_ is the killing window of DHA.

To define *E*_*PM*_, models incorporating the Loewe additivity concept (as PQ and MQ have similar modes of action) were used, which include only one parameter for the effect of the interaction between PQ and MQ (31, 32, 40). These models are more specified to the framework of drug interaction, in contrast to the statistical models that usually have multiple parameters (41, 42, 43). A detailed description of the examined models is provided in Supplementary Material 2. The final model selected was a combination of the models described in Tallarida 2006 (32) and Machado, Robinson 1994 (40):

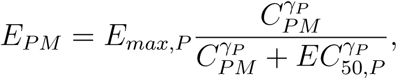

where

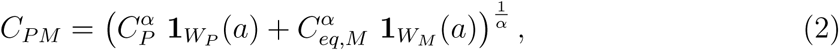

and

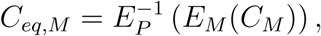

where *E*_*M*_ is the killing effect of MQ and 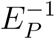 is the inverse of the killing effect of PQ, given by

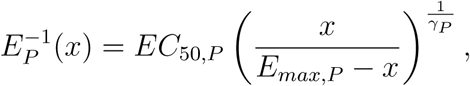

where *E*_*max*,P_ and *EC*_50,*P*_ are the maximum killing effect of PQ and the concentration at which half of the maximum killing effect is produced, respectively; *W*_*P*_ and *W*_*M*_ are the killing windows of PQ and MQ, respectively. Zero-interaction is produced by Eqn. (2) when *α* = 1; the values of 1 < *α* < ∞ and 0 < *α* < 1 produce antagonism and synergism, respectively. Note that PQ is considered to be more potent than MQ; see Supplementary Material 2 for further information.

Isobolograms, widely used in pharmacology and toxicology studies, can inform on the nature of drug-drug interactions. These present data on the parasiticidal effect of paired drug concentrations. The combination of drug concentrations is then compared with the zero-interaction isobole (also known as linear isobole) (44); see Fig. 5a. When the pairs of drugs concentrations are close to the linear isobole, zero-interaction is inferred, and when they lie significantly above or below the linear isobole, antagonism or synergism can be inferred, respectively.

**Figure 5:**
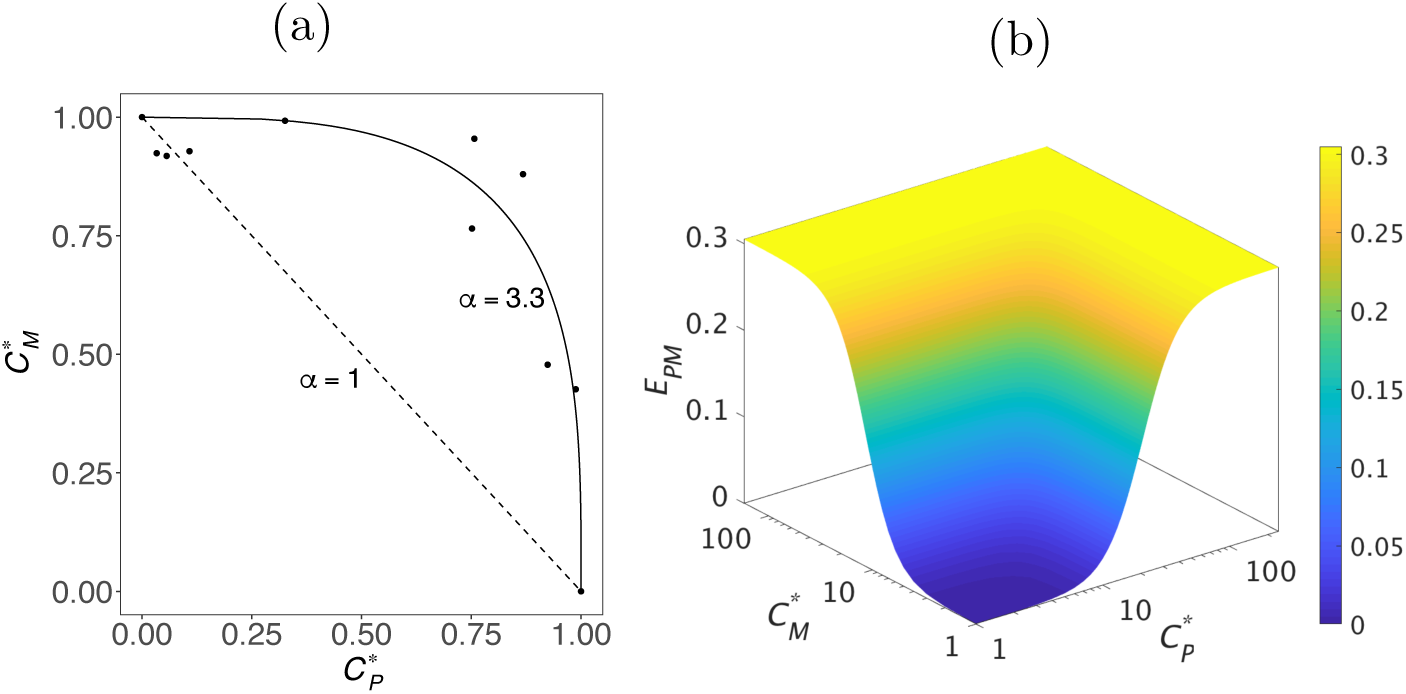
Interaction between PQ and MQ and their combined effect. (a) Isobologram presented in (13) showing an strong antagonistic interaction between PQ and MQ. The dashed and solid lines show the zero-interaction isobole and our fitted curve to the data points (estimated PQ-MQ interaction parameter is *α* = 3.3), respectively.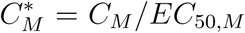 and 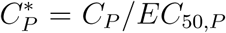 are the normalised concentrations of MQ and PQ, respectively. (b) Combined effect of PQ and MQ, *i.e. E*_*PM*_ (the 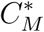 and 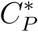 axes are log-scaled), when PQ-MQ interaction parameter (*α*) equals 3.3. The maximum killing effects and sigmoidicity of PQ and MQ are considered equal (*i.e. E*_*max,P*_ = *E*_*max,M*_ = 0.3 and *γ*_*P*_ = *γ*_*M*_ = 3) throughout the model fitting to conform with the data provided by Davis et al. (2006) (13).

Using this approach, Davis et al. 2006 (13) showed that the paired PQ and MQ data were significantly above the zero-interaction isobole (dashed line), indicating a strong antagonistic interaction between PQ and MQ; Fig. 5a. The combined killing effect of PQ-MQ, *E*_*PM*_, was fitted to this data, and PQ-MQ interaction parameter was estimated to be *α* = 3.3. Fig. 5b shows predicted *E*_*PM*_ for *α* = 3.3 for varying PQ and MQ concentrations. The killing effects of DHA and PQ-MQ were applied to Eqn. (1) to estimate the combined effect of DHA-PQ-MQ and simulate the PD model; see Supplementary Material 2 for further details.

### Model simulation

Latin Hypercube Sampling (LHS) was used to efficiently sample the parameter space (45), and simulate the PK profiles and parasitological responses. The distributions of the parameter values of the PK and PD models are presented in Tables 2 and 3, respectively. A triangular distribution was used for generating samples of *α*, with a peak at *α* = 3.3, estimated by fitting the model to the data, as explained in the previous section. The lower and upper bounds were selected to be 1 (zero-interaction) and 16 (very strong antagonism), respectively. The initial parasite burden was assumed to have a log-normal distribution with a geometric mean of 1.14 *×* 10^11^ and standard deviation of 1.13 on the log-scale; Table 3.

**Table 3:** Statistical distribution of the initial parasite burden and parameter values of the PD model.

| Parameter | Drug | Distribution | Description |
| --- | --- | --- | --- |
| $N_0$ | | $\log N(25.46, 1.13)$ | initial number of parasites |
| $\mu_{IPL}$ | | $DU(4, 16)$ | mean of initial parasites age distribution (h) |
| $\sigma_{IPL}$ | | $DU(2, 8)$ | standard deviation of initial age distribution (h) |
| $PMF$ | | $TRI(8, 12, 10)$ | parasite multiplication factor (/48 h cycle) |
| $E_{max}^*$ | DHA | $TRI(0.49, 0.69, 0.59)$ | maximum killing effect |
| | PQ | $TRI(0.19, 0.50, 0.35)$ | |
| | MQ | $TRI(0.09, 0.43, 0.26)$ | |
| $EC_{50} [ng/mL]^\dagger$ | DHA | $U(1.44, 532.05)$ | concentration producing $E_{max}/2$ effect |
| | PQ | $U(11.56, 94.19)$ | |
| | MQ | $U(20.48, 1087.22)$ | |
| $\gamma$ | DHA | $\log N(1.31, 0.65)$ | sigmoidicity of the concentration-effect curves |
| | PQ | $\log N(1.35, 0.66)$ | |
| | MQ | $\log N(0.97, 0.54)$ | |
| $\alpha$ | PQ-MQ | $TRI(1, 16, 3.3)$ | interaction parameter |

$TRI(l, h, m)$ : triangular distribution with peak at $m$ , lower limit of $l$ and higher limit of $h$ .
$DU(l, h)$ : discrete uniform distribution with lower and higher limits $l$ and $h$ , respectively.
$U(l, h)$ : continuous uniform distribution with lower and higher limits $l$ and $h$ , respectively.
$\log N(\mu, \sigma)$ : log-normal distribution derived from a normal distribution with mean $\mu$ and standard deviation $\sigma$ .
killing windows of the drugs are as follows (17): $W_D = [6 \ 44]$ , $W_P = [12 \ 36]$ and $W_M = [18 \ 40]$
\* see Supplementary Material 3 for further details
† the lower limit of the distribution of $EC_{50}$ is chosen to be the *in vitro* $IC_{50}$ of free drug, obtained by adjusting for the *in vitro* drug bindings. The higher limit is chosen to be half of the maximum drug concentration of the median of the PK profiles (17).

The probability of cure (*i.e.* 1 - probability of failure) was used as a measure of drug efficacy, and Kaplan-Meier survival analysis was carried out on the simulated parasite versus time profiles of the patients, to estimate the probabilities of cure at day 42 of follow-up. Treatment failure was defined as parasite recrudescence, in which the peripheral parasitaemia exceeded the microscopic limit of detection (50 parasite per µL or a total parasite biomass of 2.5 *×* 10^8^).

Dosing regimens recommended by WHO were used in the simulations. These included 18.0 mg/kg/day of PQ and 4.0 mg/kg/day of DHA for three days. Current guidelines recommend a total dose of 25 mg/kg MQ in combination with 4mg/kg/day of artesunate (18). Splitting the dose of MQ (8.3 mg/kg/day for three days) improves the bioavailability of MQ, is better tolerated and and has a greater efficacy (46). Higher daily doses of MQ are associated with significant side-effects (47), and thus modelling explored the minimum dosage of MQ that results in optimal efficacy. Hence, the number of days in which a 8.3 mg/kg dose of MQ was administered was varied and the corresponding TACT efficacy was estimated.

To simulate different degrees of PQ and DHA resistance, *EC*_50_, *E*_*max*_ and W were varied over the limited sampling intervals of the range of values given in Table 3. Different scenarios were considered, simulating resistance to DHA and/or PQ.

## Acknowledgments

This work was supported by the NHMRC Centres for Research Excellence in Malaria Elimination (1134989) and Infectious Diseases Modelling to Inform Public Health Policy (1078068), an NHMRC Project Grant (1100394) and an ARC Discovery Project (DP170103076). FJIF was supported by Australian Research Council Future Fellowship. RNP is a Wellcome Trust Senior Fellow in Clinical Science (200909), and JAS is a NHMRC Senior Research Fellow (1104975).

## Supplementary Materials

## 1 Other manifestations of resistance

### 1.1 Maximum killing effect, *E*_*max*_

Here we examine the probability of cure on day 42 of follow-up when *E*_*max*_ is the resistance manifestation. Fig. S1 shows the results when *E*_*max,P*_ varies across the deciles of its sampling interval; samples are taken from uniform distributions over each decile. Similar to the results of Section “Resistance to DHA”, adding one dose of MQ to the ACT (blue curve) can increase the probability of cure, but is not sufficient. In order to reach the probability of cure of above 90% for all of the deciles, we need three doses of MQ. Of interest, the magnitude of the effect of resistance on probability of cure in this case is close to that of *EC*_50_; resistance to DHA is also considered, *i.e. EC*_50,*D*_ ∈ (50, 100].

### 1.2 Killing window, *W*

We now shorten the size of killing window, *W*, of PQ for the intra-erythrocytic parasite life cycle, by increasing the lower limit of the *W* and fixing the higher limit. The results for the ACT show that shortening the killing window can significantly reduce the probability of cure, but again, adding MQ to the compound can pull up the probabilities of cure. To achieve a probability of cure of at least 90%, three 8.3 mg/kg doses of MQ are required.

**Figure S1:**
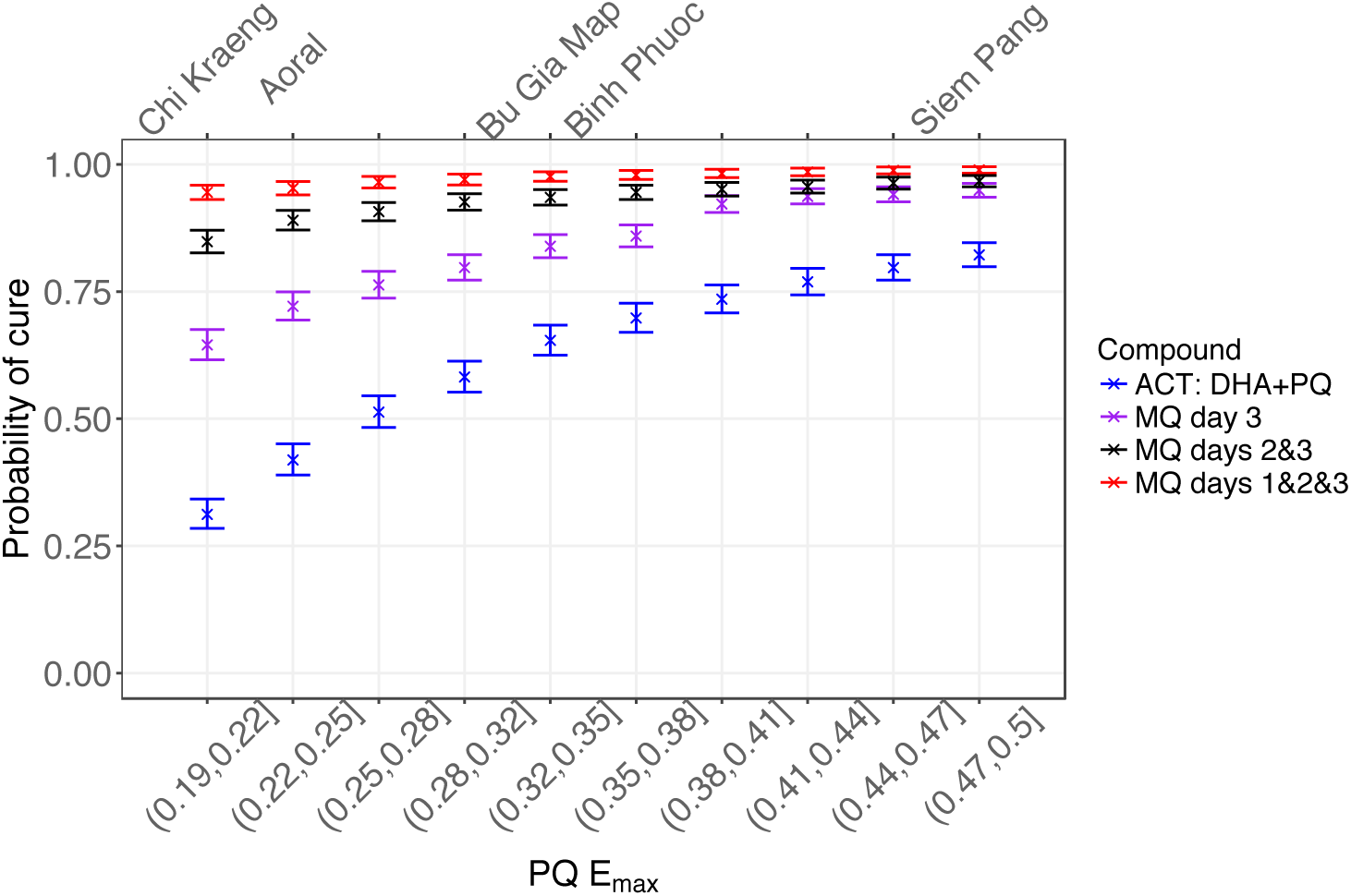
The probability of cure on day 42 of follow-up when *E*_*max*_ of PQ varies over the deciles of (0.19, 0.50]. Dosing regimens of PQ and DHA are 18.0 mg/kg and 4.0 mg/kg, respectively, on days 1, 2 and 3. Purple: a single dose of 8.3 mg/kg of MQ is added on day 3. Black: two 8.3 mg/kg doses of MQ on days 2 and 3 are added. Red: three 8.3 mg/kg doses of MQ are added on days 1, 2, 3. The top labels show the geographical regions in South-East Asia (Table 1) that have observed DHA-PQ cure rates equal to the corresponding simulated values. Error bars show the 95% confidence intervals of Kaplan-Meier analysis.

**Figure S2:**
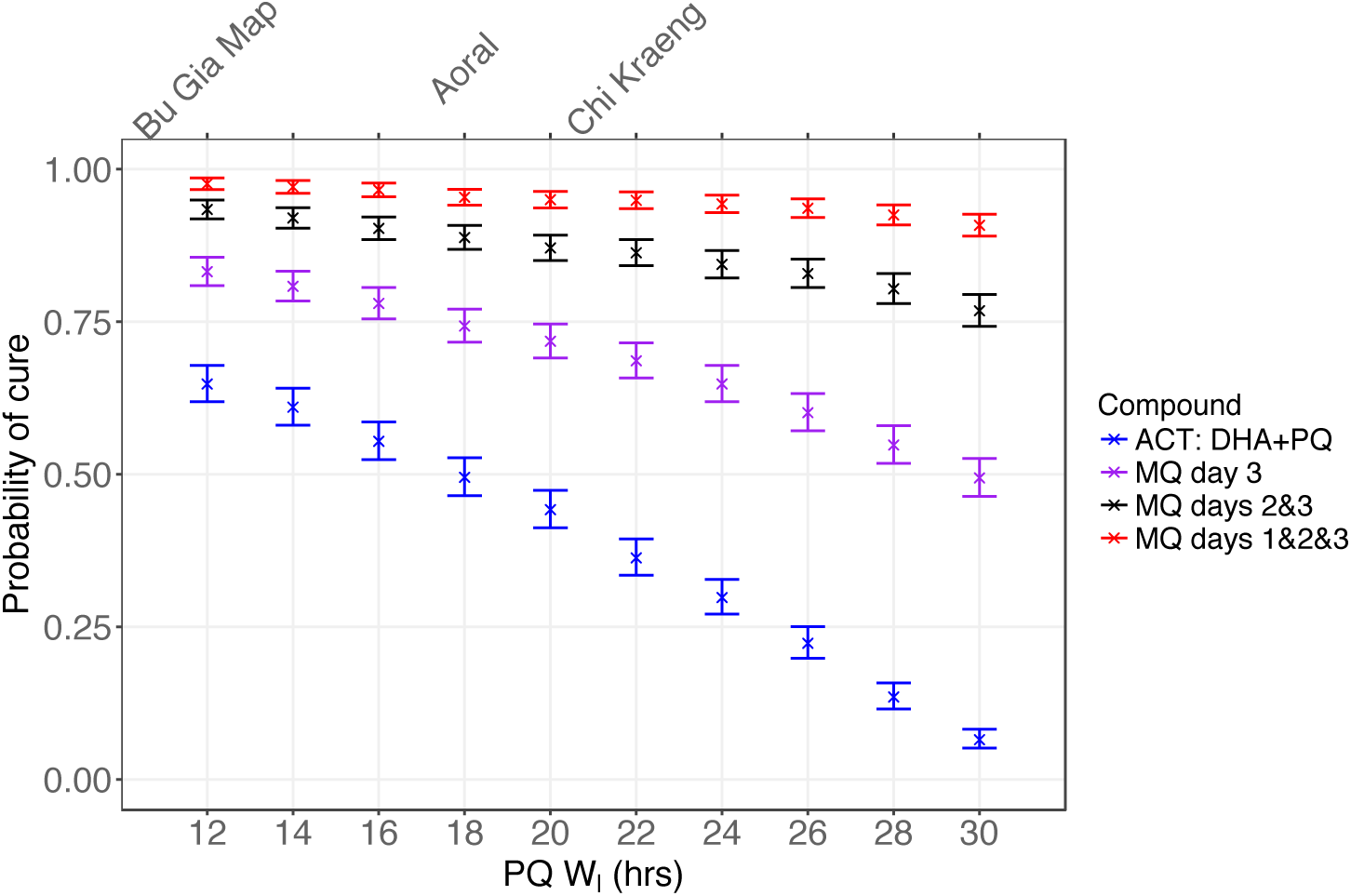
The probability of cure at day 42 of follow-up when the size of the parasite killing window *(W)* for PQ is reduced by increasing the lower limit, *W*_*l*_, from 12 to 30. The higher limit, *W*_*u*_, is constant and equal to 36 hours. Purple: a single dose of 8.3 mg/kg of MQ is added on day 3. Black: two 8.3 mg/kg doses of MQ on days 2 and 3 are added. Red: three 8.3 mg/kg doses of MQ are added on days 1, 2, 3. The top labels show the geographical regions in South-East Asia (Table 1) that have observed DHA-PQ cure rates equal to the corresponding simulated values. Error bars show the 95% confidence intervals of Kaplan-Meier analysis.

## 2 Modelling combined killing effect

### 2.1 Models of drug interaction

There are two prominent empirical approaches for modelling zero-interaction: *Loewe additivity* (34) and *Bliss independence* (35). Loewe additivity is based on the idea that two non-interacting drugs differ only in their potency, and was originally formulated as

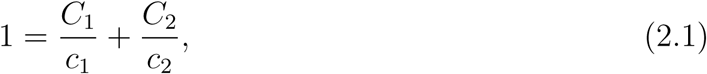

where *c*_1_ and *c*_2_ are the concentrations of drugs 1 and 2, respectively, that each individually (*i.e.* not in combination) produces a specified effect *E*_12_, and *C*_1_ and *C*_2_ are the drug concentrations in a combination that together produce *E*_12_ — for brevity, the formulae are defined for two drugs, but they can be readily extended for multiple drugs. Eqn. (2.1) is known as a *linear isobole*, which is widely used in pharmacology and toxicology as a reference to identify drug interactions. Loewe first put forward this model, which was then investigated more rigorously by Berenbaum (1985) and others.

Loewe additivity is suggested to be a suitable concept for zero-interaction when the combined drugs have similar modes of action (31, 32). However, when the drugs are believed to act independently, Bliss independence is more appropriate. This model is based on a probabilistic perspective, defined as

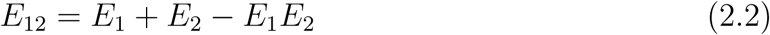

where *E*_1_ and *E*_2_ are the individually produced effects by drugs 1 and 2, respectively.

Ultimately, deviations from a selected zero-interaction reference model would determine the degree of synergistic/antagonistic interaction in certain drug combinations. Note that despite the fundamental differences of Loewe additivity and Bliss independence, it has been shown that they indicate the same nature of drug interactions in the majority of cases (48).

### 2.2 Combined effect of DHA-PQ-MQ

Statistical models can be used to define *E*_*PM*_, *e.g.* Carter et al. (1988) used a generalised linear model with the logit link function:

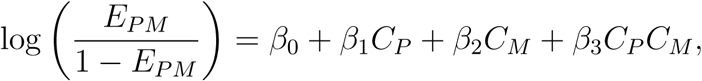

where *C*_*P*_ and *C*_*M*_ are the concentrations of PQ and MQ, respectively, and β_0_, …, β_3_ are the coefficients of the model. Similar statistical models can be found in (42, 43).

Another set of models include only one parameter to incorporate the effect of interaction (31, 40, 32). These models are more specified to the framework of drug interaction, in contrast to the statistical models. Here, we focus on the models with one parameter of interaction — noting that statistical models are shown to be readily transformable to these models, *e.g.* see (41).

One of the most frequently used models to describe the combined effect is Greco’s model (31), defined by

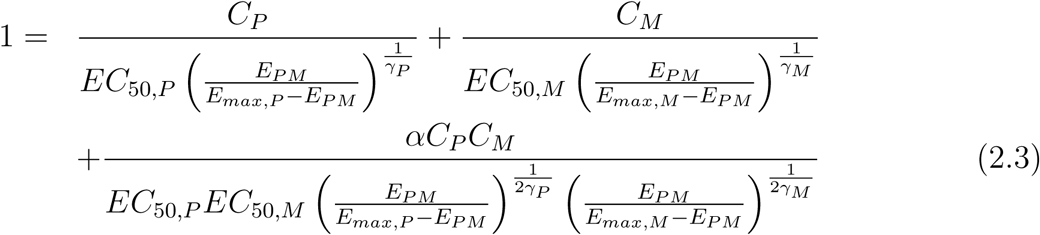

where the subscripts *P* and *M* denote which drug the parameters correspond to. The interaction parameter, *α*, incorporates the influence of the interaction between the drugs, where, for Eqn. (2.3), *α* = 0, *-*1 < *α* < 0 and *α* > 0 produce zero-interaction, antagonism and synergism, respectively. Note that we should have *E*_*PM*_ < *E*_*max,P*_ and *E*_*PM*_ < *E*_*max,M*_, otherwise, Eqn. (2.3) would not yield a real-valued solution for *E*_*PM*_. These conditions thus limit the utility of Greco’s model to cases where *E*_*max,P*_≠*E*_*max,M*_.

Tallarida (2006) put forward a broader framework based on the Loewe additivity, from which Greco’s model can be derived as a special case. In addition, it overcomes the aforementioned limitation on the values of *E*_*PM*_. In Tallarida’s approach, we first identify the more potent drug, say PQ; this can be done by carrying out *in vitro* susceptibility tests or comparing the parasite reduction ratios derived from clinical efficacy studies. Then, we find the concentration of PQ that is equally effective as MQ at concentration *C*_*M*_, using

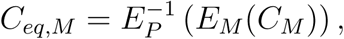

where 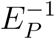 is the inverse function of *E*_*P*_, given by

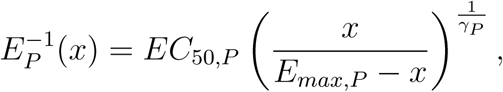

Then, the zero-interaction model is obtained via

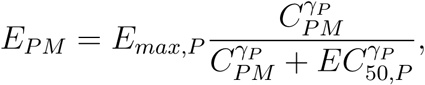

where

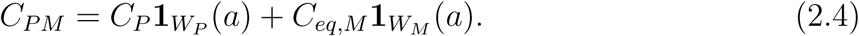

Subsequently, Eqn. (2.4) can be modified to accommodate an interaction between drugs. For example, Tallarida (2000) suggests changing this equation to *C*_*PM*_ /*α*, where*α* is the interaction parameter. However, we dismiss this method as it does not produce the observed antagonistic isoboles (see Fig. 5), hence, it will not provide a good fit to data. In order to obtain a form of *E*_*PM*_ similar to Greco’s model, Eqn. (2.3), we then modified Eqn. (2.4) to incorporate the effect of an interaction between drugs. Adding *αC*_*P*_ *C*_*eq,M*_ as an extra term to this equation provides a good fit to the data for *α* = *-*0.132, but, the resultant *E*_*PM*_ is non-onotonic, which is biologically infeasible. We also tried other terms such as 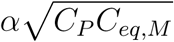 but they similarly failed to give either a good fit or a monotonic effect. Hence, the models of form Eqn. (2.3) did not produce an appropriate *E*_*PM*_, as also outlined by White et al. (2003) and Machado, Robinson (1994).

We then turned to using the model introduced by Machado, Robinson (1994):

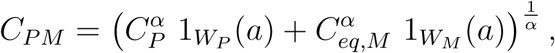

where zero-interaction is produced when *α* = 1. The values of 1 < *α* < ∞ and 0 < *α* < 1 produce antagonism and synergism, respectively. The model provides a good fit to the data (see Fig. 5a), and importantly, a biologically feasible killing effect, *E*_*PM*_ (see Fig. 5b). Therefore, we selected this model for *E*_*PM*_, and used it in the combined effect, Eqn. (1), of the TACT.

To conform with the data provided by Davis et al. (2006) (13), the maximum killing effects and sigmoidicity of PQ and MQ are considered equal (*i.e. E*_*max,P*_ = *E*_*max,M*_ = 0.3 and *γ*_P_ = *γ*_M_ = 3) throughout the model fitting. However, the considered range of variation for *α* in the simulations is significantly larger than the potential variations due to *E*_*max,P*_ ≠ *E*_*max,M*_ and/or *γ*_*P*_ ≠*γ*_*M*_, hence, these assumptions do not invalidate the results (see Table 3).

## 3 Calculating *E*_*max*_ using the parasite reduction ratio (PRR)

We are interested in finding how *E*_*max*_ is related to the parasite reduction ratio (PRR).We can estimate PRR by

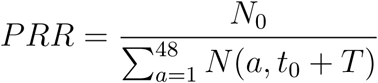

where *T* is the time when we count the number of parasites (*e.g. T* = 48 hrs) to calculate PRR, and *N*_0_ is the initial number of parasites at time *t*_0_. Then, we have

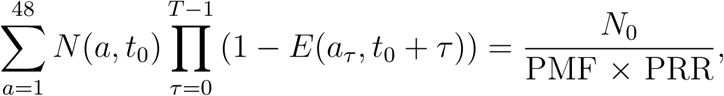

where *a*_*τ*_ = [(*a* + *τ*) mod 48]. Thus, we use numerical methods to solve the above equation for *E*_*max*_. The estimated *E*_*max*_ values are listed in Table 3. Note that it is extremely important to take account of the details of the clinical efficacy studies, by which the PRRs of the drugs are obtained. We used the following PRRs and the dosing regimens to estimate *E*_*max*_ for each drug:

- PRR_*DHA*_ = 10^4^: seven 2 mg/kg doses of DHA are administered (51).
- PRR_*PQ*_ = 2951: one 14.1 mg/kg dose of PQ is administered (52).
- PRR_*MQ*_ = 100: one 25 mg/kg dose of MQ is administered (51).

The obtained *E*_*max*_ is then used as the median of the triangular distribution (see Table 3), and the lower and higher limits of the distribution are found by

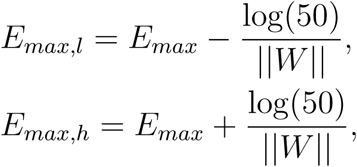

where ||*W*|| is the size of killing window of the drug (17).

